# Analysis of RNA Exosome Subunit Transcript Abundance Across Tissues: Implications for Neurological Disease Pathogenesis

**DOI:** 10.1101/2023.06.07.544082

**Authors:** Julia L. de Amorim, Don Asafu-Adjaye, Anita H. Corbett

## Abstract

Exosomopathies are a collection of rare diseases caused by mutations in genes that encode structural subunits of a ribonuclease complex termed the RNA exosome. The RNA exosome mediates both RNA processing and degradation of multiple classes of RNA. This complex is evolutionarily conserved and required for fundamental cellular functions, including rRNA processing. Recently, missense mutations in genes encoding structural subunits of the RNA exosome complex have been linked to a variety of distinct neurological diseases, many of them childhood neuronopathies with at least some cerebellar atrophy. Understanding how these missense mutations lead to the disparate clinical presentations that have been reported for this class of diseases necessitates investigation of how these specific changes alter cell-specific RNA exosome function. Although the RNA exosome complex is routinely referred to as ubiquitously expressed, little is known about the tissue- or cell-specific expression of the RNA exosome complex or any individual subunit. Here, we leverage publicly available RNA-sequencing data to analyze RNA exosome subunit transcript levels in healthy human tissues, focusing on those tissues that are impacted in exosomopathy patients described in clinical reports. This analysis provides evidence to support the characterization of the RNA exosome as ubiquitously expressed with transcript levels for the individual subunits that vary in different tissues. However, the cerebellar hemisphere and cerebellum have high levels of nearly all RNA exosome subunit transcripts. These findings could suggest that the cerebellum has a high requirement for RNA exosome function and potentially explain why cerebellar pathology is common in RNA exosomopathies.

## Introduction

Exosomopathies are a collection of rare congenital pediatric diseases resulting from missense mutations in genes encoding structural subunits of the RNA exosome complex. The RNA exosome is a ribonuclease complex required for multiple critical cellular functions that dictate gene expression and post-transcriptional regulation. One of the most well-defined fundamental roles the RNA exosome plays in gene expression is mediating the precise processing of ribosomal RNA (rRNA) required to produce mature ribosomes [1, 2]. In additional to the maturation of rRNA, the RNA exosome targets many classes of RNAs for processing, degradation, and turnover [3].

The RNA exosome complex comprises ten subunits: nine structural noncatalytic subunits and one 3’-5’ exo/endoribonuclease subunit (**Figure 1A**, PDB 6H25 [4]). Three subunits make up the cap: EXOSC1, EXOSC2, and EXOSC3; while six subunits make up the hexameric core: EXOSC4, EXOSC5, EXOSC6, EXOSC7, EXOSC8, and EXOSC9. The catalytic ribonuclease DIS3 is located at the base of the complex. The cap and core assemble to form a channel through which RNA is threaded in a 3’-5’ orientation to reach the catalytic subunit [5]. Studies in various model systems have shown that the RNA exosome complex is essential for viability [1, 6-12] and the complex is routinely referred to as ubiquitously expressed [13]. To date, six of the nine structural subunit genes that make up the RNA exosome have been linked to conditions which each involve at least some degree of cerebellar atrophy [9, 11, 12, 14-19]. Why mutations in genes that encode structural subunits of the ubiquitously expressed RNA exosome complex give rise to neurological disease, which often impacts the cerebellum, is not well understood.

**Figure 1.**
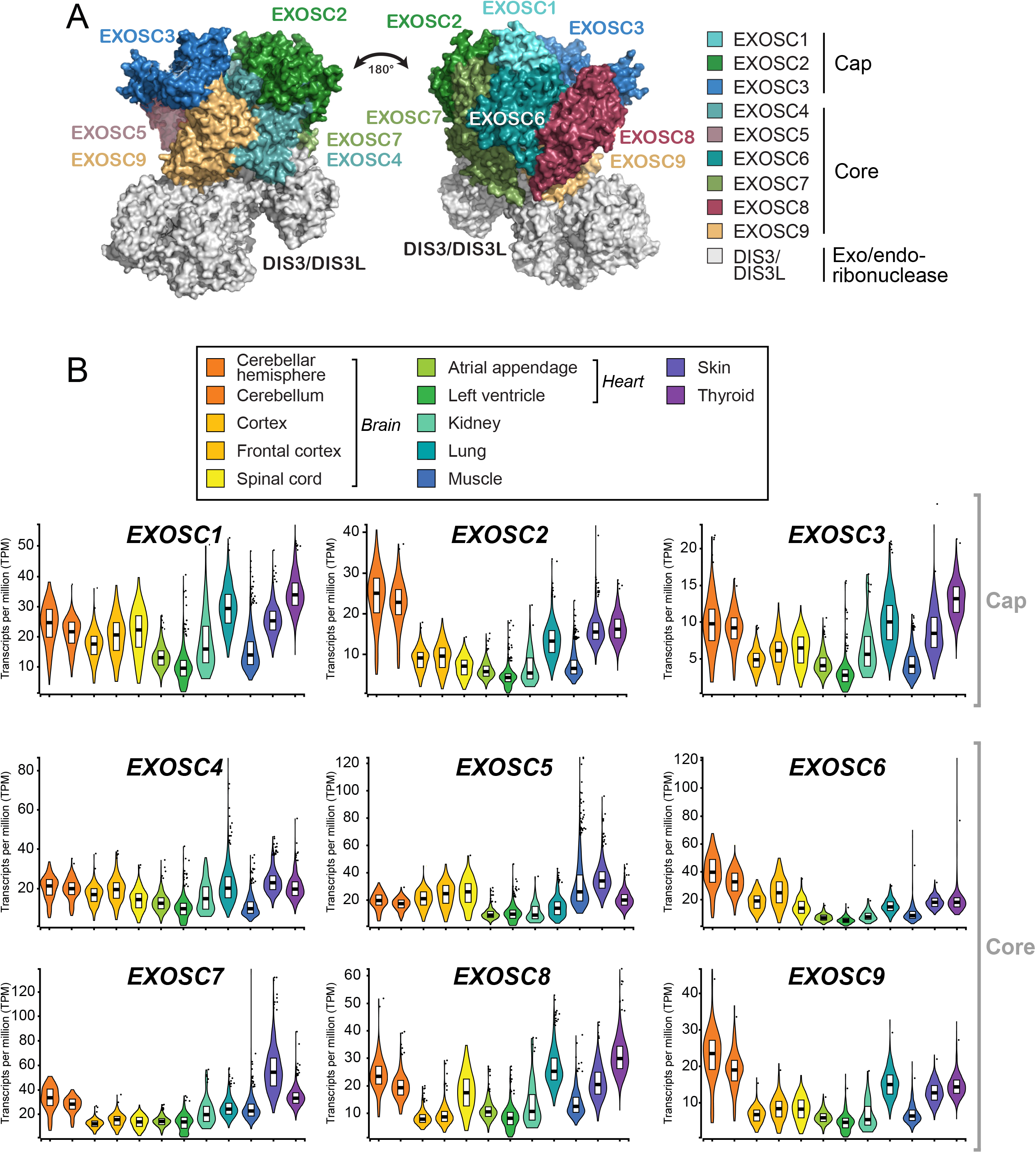
(A) The RNA exosome is a ten-subunit complex that targets multiple classes of RNA. Nine of the ten subunits of the RNA exosome are structural and nonenzymatic. Three subunits compose the cap: EXOSC1, EXOSC2, EXOSC3, and six subunits compose the hexameric core: EXOSC4, EXOSC5, EXOSC6, EXOSC7, EXOSC8, EXOSC9. The base subunit, DIS3 or DIS3L, has catalytic ribonucleolytic activity. The image is adapted from the PDB structure 6H25 [4]. (B) The nine structural subunits of the RNA exosome are expressed in all tissues examined. Transcript levels of each subunit vary; however, several subunits show high levels in the cerebellar hemisphere/cerebellum. Each structural subunit is analyzed in tissues that have been previously reported in the clinical diagnoses or causes of death for exosomopathy patients. As described in detail in the Materials and Methods, tissues examined for each RNA exosome subunit include the brain (cerebellar hemisphere, cerebellum, cortex, frontal cortex, and spinal cord), heart (atrial appendage and left ventricle), kidney, lung, muscle, skin, and thyroid. RNA levels of subunits are presented in violin plots by median transcript per million (TPM) and vary from tissue to tissue.

### The RNA exosome complex plays critical roles in gene expression in subcellular compartments

The RNA exosome localizes to both the nucleus and the cytoplasm and regulates many classes of RNA within these compartments. For example, in the nucleus, the RNA is critical for precise processing of ribosomal RNA to produce mature rRNA required for ribosomes [1, 2]. In addition, the nuclear RNA exosome targets RNA-DNA hybrids (R-loops), antisense RNAs, and small noncoding RNAs (snRNAs) for processing and/or degradation [20-23]. In the cytoplasm, the RNA exosome targets mRNA for regulatory turnover, aberrant mRNAs for decay, such as mRNAs lacking a stop codon, and double stranded RNAs (dsRNAs) as a mechanism for viral defense [24-26].

The RNA exosome is hypothesized to target specific RNAs via interactions with proteins termed cofactors [27]. In the nucleus, the RNA exosome associates with cofactors including the helicase MTR4, the TRAMP polyadenylation complex, and the MPP6 docking protein [28-30]. Cytoplasmic RNA exosome cofactors include the rRNA channeling SKI complex [31]. Given these critical interactions with the RNA exosome complex, any changes that alter the composition or conformation of the complex could have consequences for critical protein-protein interactions. Many important studies have provided insight into the function of the RNA exosome, including elegant structural studies [4, 28, 31-35], identification of key RNA substrates [3, 20, 22, 24, 36], and mechanistic insight into cofactor interactions [3, 28, 31, 37, 38]. Many studies are performed using elegant biochemical approaches with reconstituted complexes, using genetic model organisms, or in cultured cells. These studies further unearth key questions about the requirements for RNA exosome function in specific cell types and tissues in multicellular organisms. These questions have been brought into sharp focus by recent studies linking mutations in genes encoding structural subunits of the RNA exosome to human diseases, which often have neurological involvement [11].

### Clinical phenotypes of exosomopathies include neurological defects

Missense mutations in genes encoding structural subunits of the RNA exosome give rise to a class of diseases termed exosomopathies. All exosomopathies described thus far include at least one missense *EXOSC* variant [12]. Some patients are homozygous for the same missense variant, others are heterozygous for different missense variants, and some patients have a missense variant inherited in trans to a deletion [11]. Additionally, exosomopathy patients have diverse clinical presentations including cerebellar atrophy, hypotonia, and respiratory difficulties [14]. As the RNA exosome is required for many key cellular processes such as gene expression and translation, mutations that cause a complete loss of function of this complex are unlikely to be compatible with life. Thus, missense mutations linked to disease are expected to not be complete loss-of-function alleles but rather hypomorphic alleles.

As mutations in multiple genes encoding structural subunits of the RNA exosome have now been linked to disease [11], a formal postulation is that any pathogenic amino acid change that disrupts protein function could trigger a decrease in the level of that subunit and consequently the entire RNA exosome complex. Indeed, studies in fibroblasts from patients with *EXOSC* mutations support this hypothesis [19]. However, if all pathological consequences were linked to loss of the RNA exosome complex, common pathology might be shared across patients with mutations in *EXOSC* genes. In contrast, a large number of distinct clinical phenotypes have been described for individuals with exosomopathies resulting from mutations in different subunit genes.

To illustrate the diversity of pathology described for exosomopathy patients, the following descriptions compile a number of the clinical diagnoses that have been identified in patients with mutations in *EXOSC* genes. One or more patients with a missense mutation in *EXOSC1* presented with hypoplastic cerebellum, cerebral atrophy, hyperextensibility of the skin, cardiomyopathy with reduced left ventricular ejection, and hypotonia [15, 39]. A single *EXOSC1* patient died due to renal failure [15]. One or more patients with a missense mutation in *EXOSC2* presented with borderline cerebellar hypoplasia, borderline intellectual disability, mild cortical and cerebellar atrophy, and hypothyroidism [16]. Patients with missense mutations in *EXOSC3* have presented with symptoms of varying severity, classified as mild, moderate, or severe [40] The severity designations of *EXOSC3* patients are based on the genotype and clinical phenotype and clearly correlate [41]. Some *EXOSC3* patients show severe hypotonia, progressive muscular atrophy, and postnatal and progressive cerebellar volume loss [40]. Autopsies of *EXOSC3* patients demonstrated loss of neurons in the cerebellum, parts of the midbrain, and the anterior spinal cord [40]. *EXOSC3* patients with mild PCH1 often survive into early puberty and reported respiratory failure in late stages of the disease, which was rarely the cause of death [40]. *EXOSC3* patients with severe PCH1 had prenatal or congenital onset of cerebellar, pontine, and midbrain degeneration, as well as presented with severe hypotonia [40]. These patients died in infancy from postnatal respiratory failure even under constant ventilation [40]. In the case of *EXOSC5* mutations, patients required breathing support [42]. One or more patients with *EXOSC5* mutations suffered from progressive hypotonia and respiratory impairment [42]. MRIs of *EXOSC5* patients revealed reduced size of cerebellar vermis, brainstem, and pons [42]. The echocardiogram of an *EXOSC5* patient showed anomalous coronary artery fistula [42]. Several individuals with missense mutations in *EXOSC8* were reported to suffer from severe muscle weakness and died of respiratory failure before two years of age [9]. MRIs of *EXOSC8* patients showed vermis hypoplasia and cortical atrophy [9]. Autopsies of *EXOSC8* patients detected profound lack of myelin in cerebral, cerebellar white matter, and in the spinal cord [9]. Patients with missense mutations in *EXOSC9* had progressively decreased strength, severe hypotonia, recurring pulmonary infections, and respiratory insufficiencies [19]. MRIs revealed progressive cerebral and cerebellar atrophy. *EXOSC9* patients showed rapid progressive muscle weakness and respiratory impairment combined with the presence of cerebellar atrophy and motor neuronopathy [19].

In contrast to mutations in genes encoding structural subunits of the RNA exosome, mutations in the *DIS3* gene, which encodes the catalytic ribonuclease [43], have been linked to multiple myeloma rather than diseases that largely affect the brain, like the exosomopathies previously described [44]. DIS3L, an alternative ribonuclease that interacts with the nine structural subunits of the RNA exosome [5], has not been causatively linked to any inherited disease. Finally, three structural subunits of the RNA exosome, EXOSC4, EXOSC6, and EXOSC7, have not yet been linked to any reported pathology. However, we predict studies will soon surface describing new patients with mutations in these three structural subunits that may result in cerebellar atrophy.

With the small number of individuals diagnosed with exosomopathies thus far [11], more similarities may be revealed. Alternatively, disease pathology may be more tightly linked to the individual *EXOSC* genes altered. In this case, the location of the pathogenic amino acid change within a specific subunit may have a significant impact on protein-protein interactions, disrupting key interactions with specific cofactors or affecting the integrity of the complex. These changes may alter the function of individual subunits or the entire complex, ultimately affecting downstream RNA targets. There may be critical RNA targets that are important for the proper function of specific cells or tissues.

In this study, we employ publicly available transcriptomic data to explore expression of individual RNA exosome subunits in various human tissues. This analysis provides support for the characterization of the RNA exosome as ubiquitously expressed and reveals there is variability in the level of transcripts for individual RNA exosome in tissues linked to clinical pathology in exosomopathies. However, the cerebellum has high levels of transcripts encoding virtually all RNA exosome subunits. This finding could suggest that cell types within the cerebellum require a high level of RNA exosome function for proper growth and/or maintenance.

## Materials and Methods

### Genotype-Tissue Expression (GTEx) project

GTEx release v8 includes whole genome sequencing (WGS) and RNA sequencing (RNA-seq) data from 54 tissues from 948 post-mortem individuals (312 females, 636 males; age 20-70). Each genotyped tissue has at least 70 samples. Violin plots and heatmap were generated to view median transcript per million (TPM) for the structural RNA exosome subunits: *EXOSC1, EXOSC2, EXOSC3, EXOSC4, EXOSC5, EXOSC6, EXOSC7, EXOSC8*, and *EXOSC9*. The number of genotyped donors for each tissue is as follows: brain/cerebellar hemisphere, n = 175; brain/cerebellum, n = 209; brain/cortex, n = 205; brain/frontal cortex, n = 175; brain/spinal cord (cervical c-1), n = 126; heart/atrial appendage, n = 372; heart/left ventricle, n = 386; kidney/cortex, n = 73; lung, n = 578; muscle/skeletal, n = 706; skin/not sun exposed (suprapubic), n = 517; thyroid, n = 574. The cerebellar hemisphere samples refer to the entire cerebellum and were preserved as frozen tissue. The cerebellum samples were procured from the right cerebellum and were preserved in a fixative. The cortex procured from the brain sampled the right cerebral pole cortex and was preserved in a fixative. The frontal cortex from the brain also sampled the right cerebral frontal pole cortex and was preserved as frozen tissue. The data used for the analyses described in this manuscript were obtained from the GTEx portal on April 20, 2023.

## Results and Discussion

### A comparative analysis of RNA exosome subunits reveals disparate requirements of transcripts in tissues

Tissues examined in this study are chosen based on clinical diagnoses or causes of death listed for individuals with exosomopathies. The varied clinical pathologies reported in RNA exosomopathies led us to include the following 12 human samples in this analysis: (1) cerebellar hemisphere/(2) cerebellum in the brain, (3) cortex/(4) frontal cortex in the brain, (5) spinal cord, (6) atrial appendage/(7) left ventricle in the heart, (8) kidney, (9) lung, (10) muscle, (11) skin, and (12) thyroid.

We examined the median expression as transcript per million (TPM) for each structural subunit of the RNA exosome in each of the tissues analyzed (**Figure 1B**). Violin plots show the levels of subunit transcripts compared between tissues. *EXOSC3* transcripts have the lowest overall TPM and *EXOSC5, EXOSC6*, and *EXOSC7* have the highest overall TPM across all tissues analyzed. Several of the RNA exosome subunit transcripts are expressed at the highest level in cerebellar hemisphere/cerebellum as compared to other tissues, including *EXOSC2, EXOSC6*, and *EXOSC9*. Importantly, the number of transcripts as reported by TPM values does not necessarily correlate with the amount of protein present in cells or tissues [45]. *EXOSC2* and *EXOSC9* patients have distinct clinical diagnoses yet both of these subunits have some of the highest transcript levels (∼20-25 median TPM) in cerebellar hemisphere/cerebellum.

To visualize the median TPM for clustered RNA exosome subunits and the tissues analyzed, we produced a heatmap using GTEx (**Figure 2A**). The cerebellar hemisphere/cerebellum, skin, and thyroid show the highest level of RNA exosome subunit transcripts. The atrial appendage and left ventricle of the heart have the lowest levels of subunit transcripts overall. *EXOSC3* transcripts are detected at the lowest levels across all tissues. To compare all RNA exosome subunits for each tissue examined, we mapped the log_10_ of the median TPM in violin plots (**Figure 2B**). Observations made from the heatmap are supported by the data provided in the violin plots.

**Figure 2.**
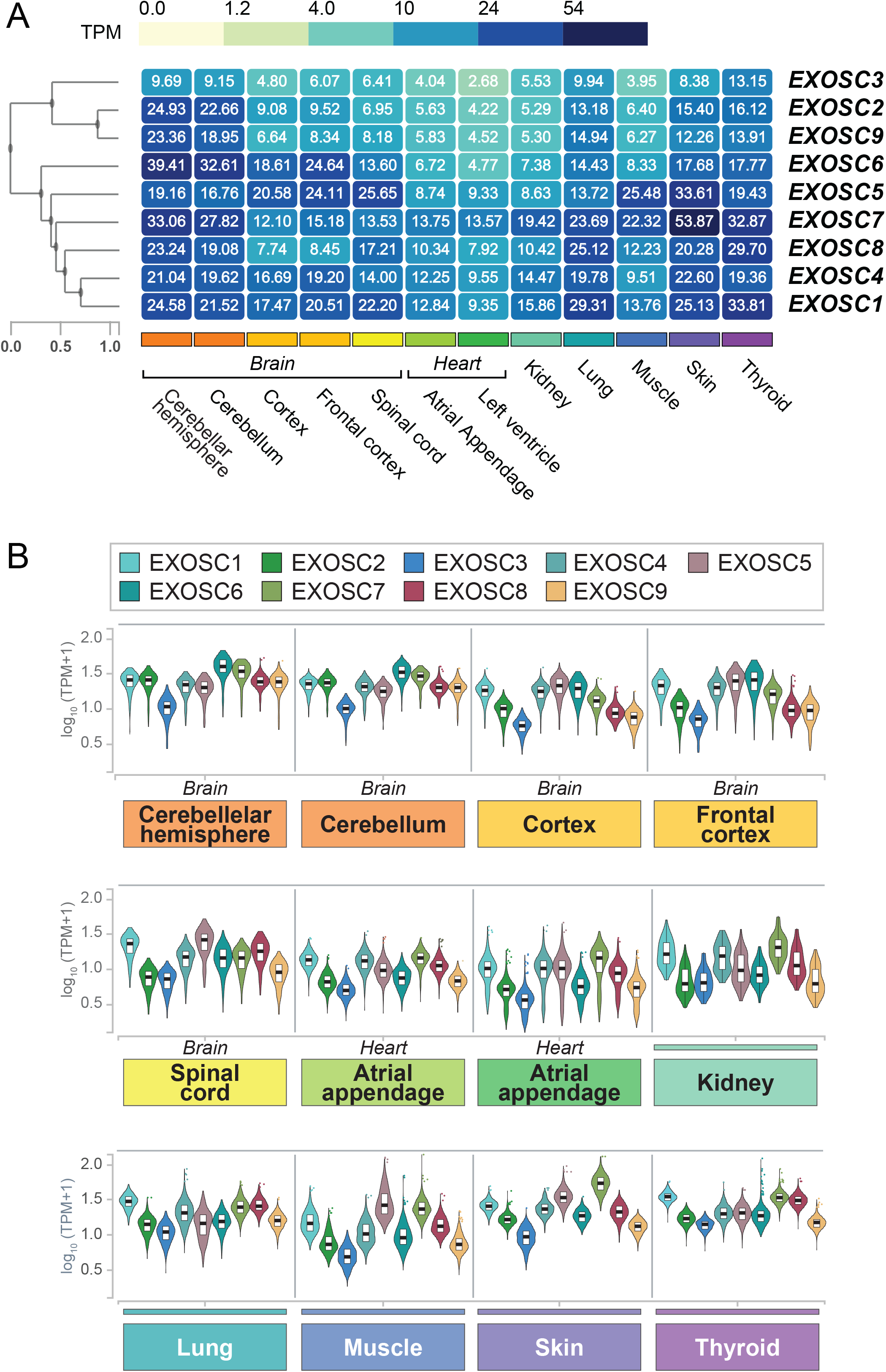
(A) RNA exosome subunit transcript levels across a set of tissues implicated in clinical diagnoses or causes of death in exosomopathy patients are shown in a heatmap. The data generated provide visualization of transcript levels across tissues as illustrated by median TPM and are clustered by RNA exosome subunit. (B) A comparative analysis of subunits reveals variable transcript levels across different tissues. Median TPM is adjusted by log_10_ TPM for comparison.

We leveraged the GTEx consortium to visualize the transcript levels of RNA exosome subunits in brain regions, heart regions, kidney, lung, muscle, skin, and thyroid by median TPM in violin plots and heatmaps. These data show that overall, the tissues representing the cerebellum have high levels of RNA exosome subunit transcripts across the various structural subunits. Specifically, the cerebellar hemisphere/cerebellum has the highest levels of *EXOSC2, EXOSC6*, and *EXOSC9*. Skin has the highest levels of *EXOSC4, EXOSC5*, and *EXOSC7*. Thyroid has the highest levels of *EXOSC1, EXOSC3*, and *EXOSC8*. The left ventricle of the heart has the lowest transcript levels of *EXOSC1, EXOSC2, EXOSC3, EXOSC6*, and *EXOSC9*. The brain cortex has the lowest transcript levels of *EXOSC7* and *EXOSC8*. Muscle has the lowest levels of *EXOSC4*, and kidney has the lowest levels of *EXOSC5*. Additionally, these data show that *EXOSC1* has the highest transcript levels in the lung and thyroid. *EXOSC5* has the highest transcript levels in the cortex, spinal cord, and muscle. *EXOSC6* has the highest transcript levels in the cerebellar hemisphere/cerebellum and frontal cortex. *EXOSC7* has the highest transcript levels in the atrial appendage and left ventricle of the heart, kidney, and skin. *EXOSC3* shows the lowest transcript level in all tissues examined except the kidney; however, levels in the kidney are relatively low compared with other subunit transcripts (*EXOSC1, EXOSC4, EXOSC5, EXOSC6, EXOSC7*, and *EXOSC8*). Levels of *EXOSC2* transcript are lowest in kidney.

Across nearly all tissues examined, *EXOSC3* transcript levels are detected at the lowest levels. The kidney is the exception and comparatively *EXOSC3* levels are only slightly above the lowest (lowest TPM = 5.29, highest TPM = 19.42, *EXOSC3* TPM = 5.53). Although little is known about the assembly of the RNA exosome complex *in vivo*, there could be a limiting subunit that determines overall levels of the complex. The observation that the *EXOSC3* transcript is low in all tissues examined could suggest that EXOSC3 levels are limiting for RNA exosome complex assembly or steady-state levels. Interestingly, mutations in the *EXOSC3* gene were the first identified and linked to human disease [46]. Perhaps, if levels of EXOSC3 are limiting, even small changes that alter the function or level of EXOSC3 could cause pathology. Indeed, the most mutations have now been identified and reported in *EXOSC3* [41], which could be consistent with this RNA exosome subunit being most vulnerable to even minor changes in function or levels. Alternatively, as EXOSC3 was the first subunit of the RNA exosome linked to disease, the larger number of cases could simply represent ascertainment bias. As additional exosomopathy cases are identified and described, differences in the numbers of patients and types of pathologies associated with mutations in different *EXOSC* genes will likely be clarified. The results reported here suggest that some tissues, such as the cerebellar hemisphere/cerebellum, skin, and thyroid may require more RNA exosome function as compared to other tissues such as the heart, brain cortex, muscle, and kidney. The data presented here may explain why exosomopathy patients with missense mutations in structural RNA exosome subunit genes have the most well-defined phenotypes in the cerebellum as compared to other tissues. However, steady-state transcript levels often do not directly translate to protein levels [45] as there are many additional regulatory steps that determine steady-state protein levels. Further proteomic analysis of RNA exosome subunits in specific cell types, such as those that are abundant in the cerebellum, could provide insight into why mutations in genes encoding structural subunits of the RNA exosome often cause cerebellar pathology. RNA exosome function is determined by many protein-protein interactions with cofactors. The cell-specificity and/or cell-specific interactions of well-defined RNA exosome cofactors has not yet been determined and novel interactors are continuously discovered [23]. Changes in cell-specific interactions could explain the phenotypes observed in exosomopathy patients.

This study provides insight into the expression of various structural subunits of the RNA exosome in human tissues, supporting the common statement that the RNA exosome complex is ubiquitously expressed. This work reveals high transcript levels for multiple RNA exosome subunits in cerebellar samples, which could begin to explain why exosomopathies often present with cerebellar involvement.

## Acknowledgements

Data provided by the Genotype-Tissue Expression (GTEx) project was used for the analyses described in this manuscript and were obtained on April 20, 2023. We thank members of the Corbett lab for valuable feedback and Dr. Elizabeth Leslie for discussion regarding disease prevalence. We thank our collaborators including Ambro van Hoof. This work was supported by the National Institutes of General Medical Sciences R01-GM130147 to AHC and AvH.

## List of abbreviations

RNA: ribonucleic acid
EXOSC: RNA exosome component
PCH: pontocerebellar hypoplasia
GTEx: Genotype-Tissue Expression project
RNA-seq: RNA-sequencing
WGS: Whole genome sequencing
TPM: transcript per million
TRAMP: Trf4/5, Air1/2, Mtr4 Polyadenylation complex

